# Dietary Sodium Deprivation Remodels the Serum Lipidome and Reveals Systemic Metabolic Adaptation in Rats

**DOI:** 10.64898/2026.06.26.734806

**Authors:** Joshua Cornman-Homonoff, Saravanan Kolandaivelu, Jennifer Veverka, Justin T. Kupec, Geoffrey I. Sandle, Vazhaikkurichi M. Rajendran

## Abstract

**Background:** Dietary sodium restriction is a common nutritional and physiological challenge that activates electrolyte-conserving endocrine pathways, but its impact on systemic lipid metabolism remains incompletely defined. We examined whether short-term dietary sodium deprivation alters the circulating lipidome and identifies lipid signatures of metabolic adaptation.

**Methods:** Male Sprague-Dawley rats were maintained on sodium-sufficient (NaS) or sodium-deprived (NaD) diets for 7 days (n=3 per group). Serum lipids were profiled by untargeted LC-MS/MS in positive and negative ion modes. Lipidomic differences were evaluated using class-level and species-level analyses, principal component analysis, volcano plots, heatmaps, and pathway-oriented interpretation.

**Results:** NaD rats exhibited a distinct serum lipidomic profile compared with NaS controls, indicating global remodeling of circulating lipid composition. Sodium deprivation produced class-specific and species-resolved changes, including selective depletion of subsets of neutral lipid species, prominent wax ester remodeling, increased phosphatidylcholine and lysophosphatidylcholine abundance, and altered acylcarnitine profiles. These signatures are consistent with coordinated changes in lipid storage, membrane phospholipid turnover, and mitochondrial fatty-acid handling.

**Conclusions:** Dietary sodium deprivation induces coordinated serum lipidome remodeling in rats, supporting the concept that nutritional electrolyte status can influence systemic lipid metabolism. These exploratory findings identify sodium deprivation as a metabolic stressor linked to neutral lipid mobilization, phospholipid remodeling, and altered mitochondrial substrate handling, and provide a foundation for future mechanistic studies.

## BACKGROUND

Dietary sodium intake is a major nutritional determinant of extracellular fluid balance, epithelial ion transport, and neurohormonal regulation [1, 2]. Sodium restriction activates the renin-angiotensin-aldosterone system (RAAS), which coordinates adaptive responses to preserve sodium balance [3, 4]. Although these pathways have been studied extensively in renal, cardiovascular, and intestinal physiology, less is known about whether dietary sodium availability also influences systemic metabolic organization.

Metabolic homeostasis depends on coordinated lipid pathways that govern energy storage, membrane composition, lipid signaling, and mitochondrial substrate utilization [5, 6]. Circulating lipid species, including triglycerides, phospholipids, lysophospholipids, and acylcarnitines, provide integrative indicators of nutritional and metabolic state [7–9]. Advances in lipidomics now enable high-resolution profiling of these lipid classes, offering a systems-level approach to define how dietary and physiological stressors reshape metabolic networks [6, 10, 11].

Emerging evidence suggests that endocrine and ionic signals can influence metabolic pathways beyond their canonical roles in ion and fluid homeostasis [12, 13]. Aldosterone signaling has been implicated in regulation of mitochondrial function and cellular metabolism, raising the possibility that sodium imbalance may drive coordinated lipid metabolic reprogramming [14–16]. However, the impact of dietary sodium deprivation on systemic lipid organization and metabolic adaptation has not been systematically defined. Here, we used untargeted serum lipidomics to examine how short-term sodium deprivation reshapes the circulating lipidome. Our findings reveal coordinated remodeling of neutral lipids, phospholipids, and acylcarnitines, supporting dietary sodium deprivation as a nutritional-electrolyte stress that influences systemic lipid metabolism.

## MATERIALS AND METHODS

### Animals and dietary intervention

Male Sprague-Dawley rats (125-150 g; Charles River Laboratories) were housed under standard laboratory conditions with controlled temperature and a 12-hour light/dark cycle. Animals were randomly assigned to either a sodium-sufficient (NaS) or sodium-deprived (NaD) diet for 7 days. The NaD diet (MP Biomedicals, catalog #0296023210) contained negligible sodium content, while NaS control animals received standard chow. This dietary design was used to model short-term nutritional sodium deprivation. All procedures were conducted in accordance with institutional guidelines for animal care and use.

### Serum collection

At the end of the dietary intervention, animals were euthanized, and blood samples were collected by cardiac puncture. Serum was isolated by centrifugation at 4°C and stored at −80°C until lipidomic analysis.

### Lipid extraction and LC–MS/MS analysis

Serum lipid analyses were commercially performed by Creative Proteomics (Shirley, NY, USA). In brief, serum lipids were analyzed by untargeted liquid chromatography–tandem mass spectrometry (LC–MS/MS) in both positive and negative ion modes. Lipids were extracted from serum using organic solvent–based methods. Chromatographic separation was achieved by reverse-phase liquid chromatography, followed by detection on a high-resolution mass spectrometer. Lipid species were annotated based on accurate mass, retention time, and MS/MS fragmentation patterns using established lipidomics databases and annotation pipelines. Data acquisition in both positive and negative ionization modes was used to maximize lipid-class coverage.

### Data processing and normalization

Raw peak intensities were extracted and normalized to total ion intensity for each sample to account for technical variability. Lipid species with missing values across replicates were excluded from downstream analyses. For each lipid species, mean abundance values were calculated from biological replicates (NaS: N1–N3; NaD: A1–A3). Fold change was calculated as the ratio of mean NaD to mean NaS abundance, and log₂-transformed values were used for visualization and statistical analyses.

### Statistical analysis

Statistical comparisons between NaS and NaD groups were performed using unpaired two-tailed Welch’s t-tests. Given the exploratory design and small sample size (n=3 per group), statistical testing was used to prioritize lipid features and support class-level trends; interpretation emphasized effect size, directionality, and concordance across related lipid species/classes in addition to nominal P values. For visualization, -log10-transformed P values were used in volcano plots. Principal component analysis (PCA) was performed on log-transformed, standardized lipid intensity values to assess global lipidomic differences between groups. Processed positive- and negative-ion mode lipidomics feature tables, including lipid annotations, replicate peak areas, group means, fold changes, and statistical outputs, are provided as Supplementary Tables ST1 and ST2.

### Lipid class and species-level analyses

For class-level analyses, lipid species were grouped by lipid class (e.g., TG, CE, WE, PC, LPC, AcCa), and class-summed abundance values were calculated by summing individual species intensities within each class. Species-level analyses were performed to identify the most differentially regulated lipids based on fold change and statistical significance. Both class-level and species-level analyses were used to distinguish global from selective lipid remodeling.

### Heatmap and clustering analysis

Heatmaps were generated using the top differentially regulated lipid species ranked by P value and/or effect size. Data were log-transformed and row-normalized (z-score) prior to visualization. Hierarchical clustering was applied to identify patterns of lipid regulation across samples.

### Pathway-oriented interpretation

Functional pathway interpretation was performed using MetaboAnalyst 6.0 based on lipid class changes. Pathways related to fatty-acid metabolism, phospholipid remodeling, and mitochondrial substrate handling were considered to aid biological interpretation of lipidomic alterations.

## RESULTS

Dietary sodium deprivation induces global serum lipidomic reorganization: Untargeted lipidomic profiling revealed a robust reorganization of the circulating lipidome under sodium deprivation (Fig. 1). Principal component analysis demonstrated clear separation between NaS and NaD groups, indicating a systemic shift in lipid composition (Fig. 1A). Differential abundance analysis confirmed widespread remodeling across lipid species, and clustering analysis revealed a reproducible lipidomic signature associated with dietary sodium deprivation (Figs. 1B and 1C).

**Figure 1.**
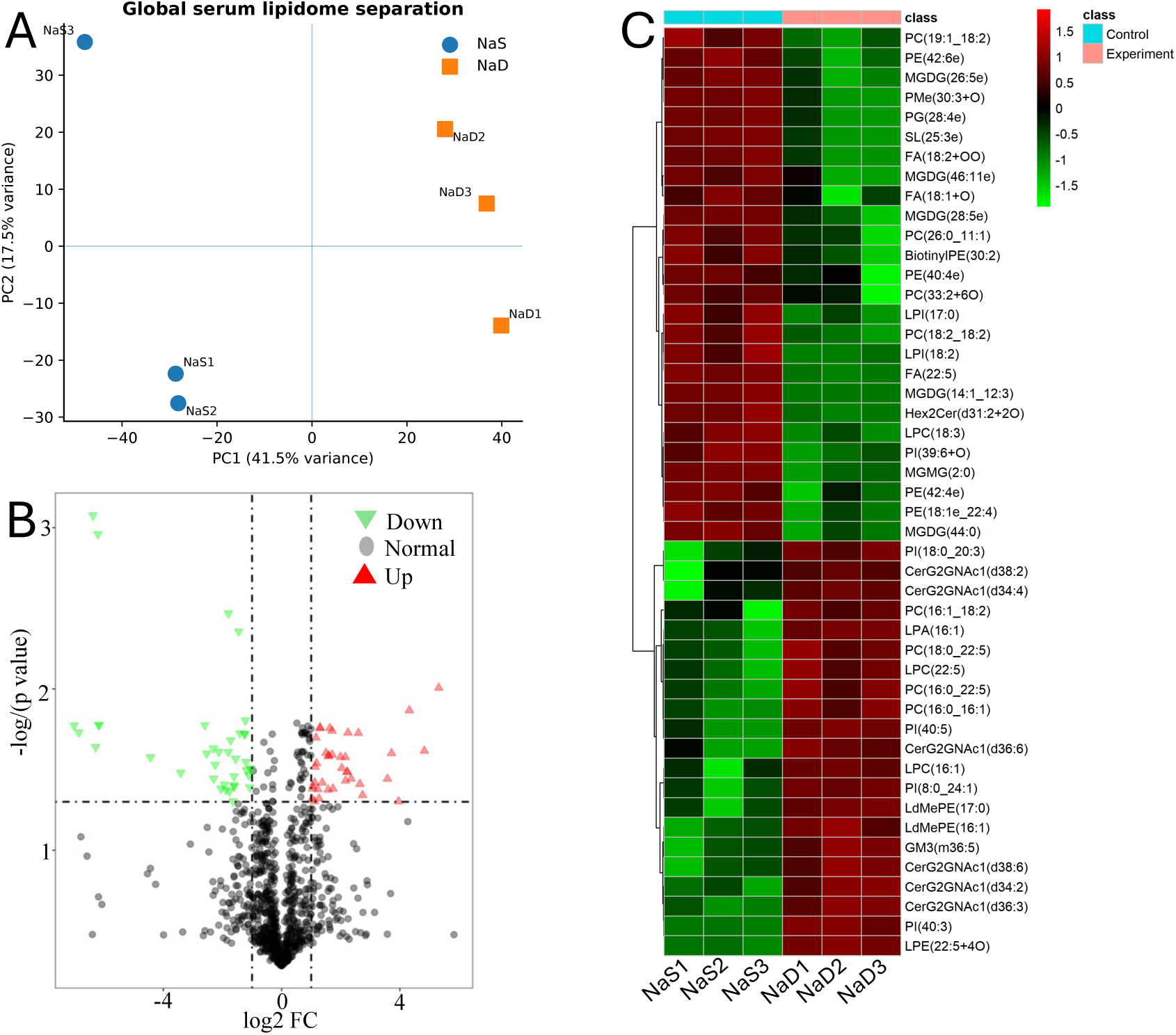
Dietary sodium deprivation remodels the serum lipidome. (A) Principal component analysis (PCA) of untargeted lipidomics data demonstrates clear separation between sodium-sufficient (NaS) and sodium-deprived (NaD) rats, indicating a coordinated shift in the circulating lipidome (n=3 per group). (B) Volcano plot showing differential serum lipid species between NaD and NaS conditions. Each point represents a lipid species; the x-axis indicates log2 fold change (NaD/NaS), and the y-axis shows -log10(P value) from two-tailed unpaired comparisons. Dashed lines denote the no-change axis and significance/effect-size thresholds used to highlight differentially altered lipids. (C) Heatmap of the top differentially altered lipid species, displayed as row z-scores across individual biological replicates (NaS1-NaS3, NaD1-NaD3). Together, these analyses show that dietary sodium deprivation drives structured remodeling of circulating lipid composition rather than isolated changes in a few metabolites.

Neutral lipid remodeling suggests selective mobilization of lipid reserves: Analysis of neutral lipid classes revealed that sodium deprivation did not induce uniform depletion of triglycerides or cholesteryl esters at the class level (Fig. 2A and Supplementary Fig. S1). Instead, species-level analysis demonstrated selective depletion of subsets of these lipids, particularly within the wax ester pool (Fig. 2B). These findings indicate that dietary sodium deprivation elicits targeted mobilization of specific lipid reserves rather than generalized lipid loss, suggesting a regulated metabolic adaptation to nutritional-electrolyte stress.

**Figure 2.**
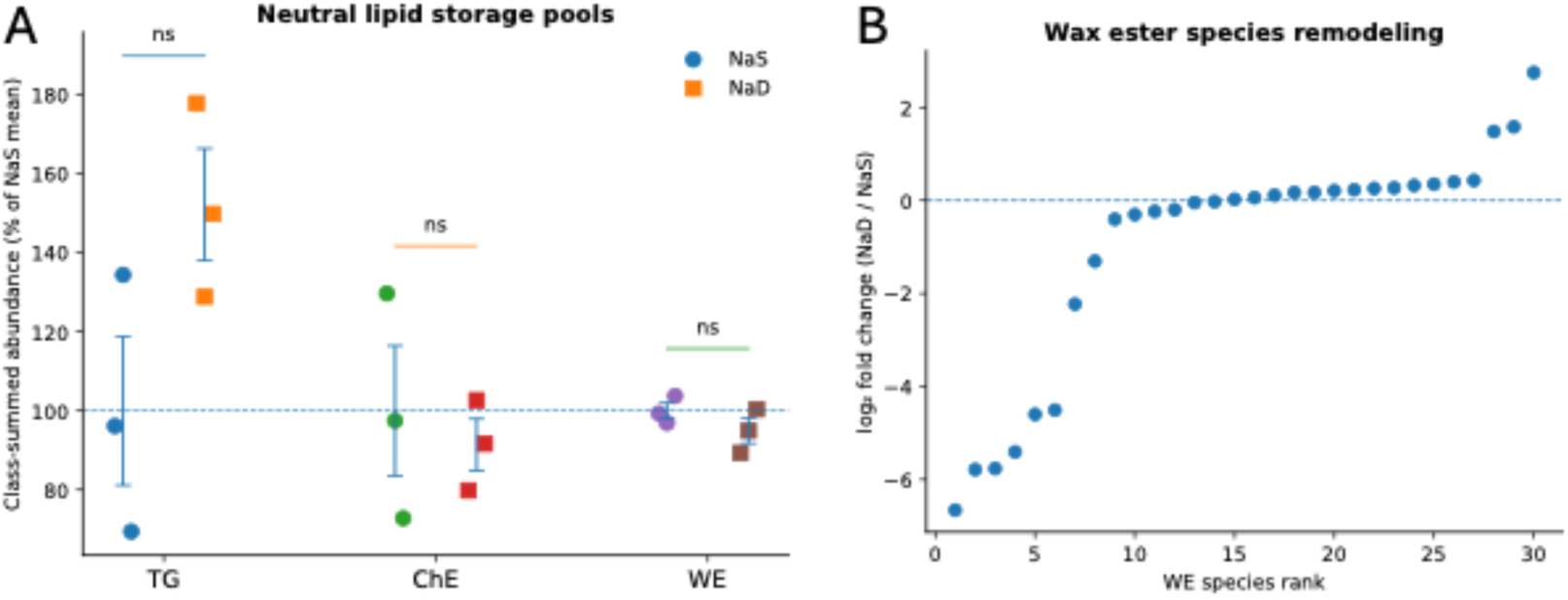
Dietary sodium deprivation remodels neutral lipid storage pools and profoundly depletes wax esters. (A) Class-summed abundances of major neutral lipid storage classes-triglycerides (TG), cholesteryl esters (CE), and wax esters (WE)-in serum from sodium-sufficient (NaS) and sodium-deprived (NaD) rats. Values are normalized to the NaS group mean (NaS=100%, dashed line) and shown as individual biological replicates (n=3 per group) with mean ± SEM. “ns” indicates not significant by two-tailed unpaired t-test. (B) Species-level remodeling of WE shown as log2 fold change (NaD/NaS) for all detected WE species, ordered by fold change; the dashed line indicates no change (log2FC=0). Together, these panels demonstrate marked depletion of WE species under sodium deprivation and class-level restructuring of neutral lipid storage pools.

Phospholipid remodeling suggests altered membrane dynamics: Sodium deprivation was associated with increased abundance of phosphatidylcholine and lysophosphatidylcholine species (Figs. 3A and 3B). Given the central role of these lipids in membrane structure and signaling [17, 18], these changes suggest enhanced phospholipid turnover and remodeling of cellular membranes. Species-level analysis supported coordinated reorganization of phospholipid composition.

**Figure 3.**
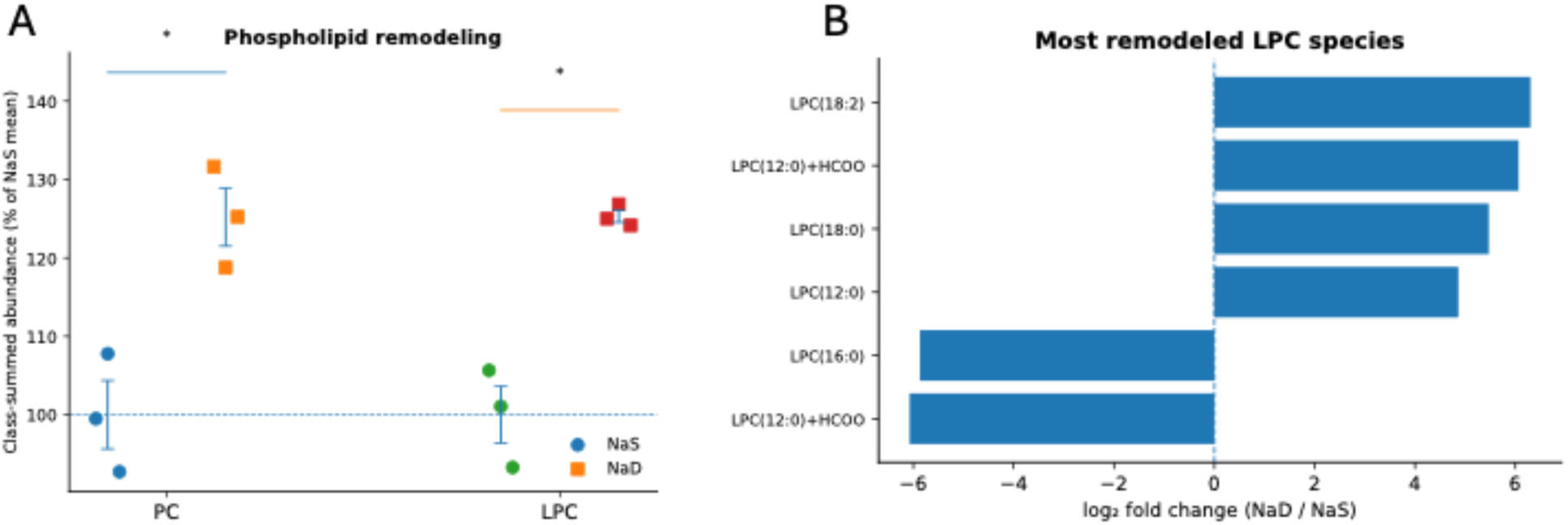
Dietary sodium deprivation drives phospholipid remodeling and selectively alters lysophosphatidylcholine species. (A) Class-summed abundances of the major membrane phospholipid classes phosphatidylcholine (PC) and lysophosphatidylcholine (LPC) in serum from sodium-sufficient (NaS) and sodium-deprived (NaD) rats. Values are normalized to the NaS group mean (NaS=100%, dashed line) and shown as individual biological replicates (n=3 per group) with mean ± SEM. Significance was assessed using a two-tailed unpaired t-test; P<0.05. (B) Bar plot of the most remodeled LPC species, shown as log2 fold change (NaD/NaS). Positive values indicate enrichment under sodium deprivation, whereas negative values indicate depletion. Together, these data indicate coordinated phospholipid remodeling with species-specific shifts in LPC composition under sodium deprivation.

Acylcarnitine remodeling suggests altered mitochondrial fatty-acid handling: Acylcarnitine profiling revealed increased total abundance alongside heterogeneous species-level changes (Fig. 4). Because acylcarnitines serve as intermediates in mitochondrial fatty-acid metabolism [19, 20], these findings suggest reorganization of substrate flux rather than uniform activation of beta-oxidation. This pattern is consistent with adaptive metabolic reprogramming at the level of mitochondrial fatty-acid handling.

**Figure 4.**
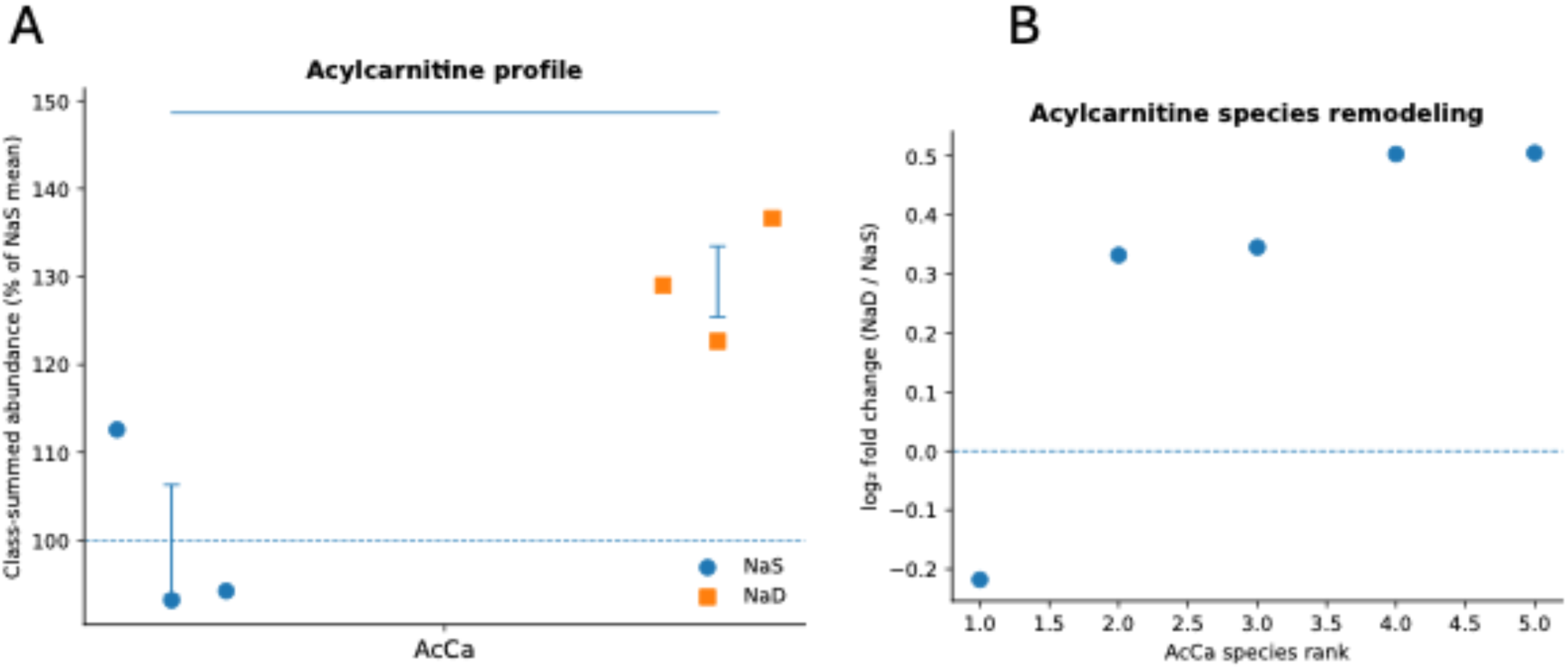
Dietary sodium deprivation increases circulating acylcarnitines and remodels acylcarnitine species composition. (A) Class-summed abundance of acylcarnitines (AcCa) in serum from sodium-sufficient (NaS) and sodium-deprived (NaD) rats. Values are normalized to the NaS group mean (NaS=100%, dashed line) and shown as individual biological replicates (n=3 per group) with mean ± SEM. (B) Species-level remodeling of acylcarnitines shown as log2 fold change (NaD/NaS) for detected AcCa species, plotted by species rank (ordered by fold change); the dashed line indicates no change (log2FC=0). Together, these panels indicate coordinated enrichment of circulating acylcarnitines under sodium deprivation, consistent with altered mitochondrial fatty-acid flux.

## DISCUSSION

This study demonstrates that short-term dietary sodium deprivation induces coordinated remodeling of the circulating lipidome, revealing an underrecognized link between nutritional electrolyte status and systemic metabolic adaptation. A key feature of this response is selective remodeling of neutral lipid pools, in which subsets of triglycerides, cholesteryl esters, and wax esters are preferentially depleted (Fig. 2 and Supplementary Fig. S1). This pattern suggests regulated mobilization of specific lipid species rather than global lipolysis, indicating a controlled metabolic response to sodium deprivation.

In parallel, we observed phospholipid remodeling, characterized by increased phosphatidylcholine and lysophosphatidylcholine abundance (Fig. 3). These changes are consistent with enhanced membrane turnover and may reflect adaptive restructuring of cellular membranes [21, 22]. Such remodeling could influence membrane fluidity, lipid signaling, and organelle function, linking lipid composition to cellular adaptation [23, 24]. Acylcarnitine profiling further supports this interpretation, revealing heterogeneous changes that suggest altered mitochondrial substrate utilization (Fig. 4) [25]. Rather than uniformly increasing fatty-acid oxidation, sodium deprivation appears to alter the balance of metabolic intermediates, consistent with a shift in mitochondrial metabolic state.

Mechanistically, these findings are compatible with activation of endocrine electrolyte-stress pathways such as RAAS (Fig. 5). Aldosterone has been implicated in regulating mitochondrial function and cellular metabolism, providing a plausible link between sodium imbalance and lipid remodeling [16, 26]. Although direct mechanistic testing was not performed, the observed lipidomic patterns are consistent with coordinated metabolic reprogramming driven by hormonal and ionic signals. Importantly, our findings extend the concept of dietary sodium regulation beyond classical ion transport and fluid balance, positioning nutritional electrolyte status as a determinant of systemic metabolic organization.

**Figure 5.**
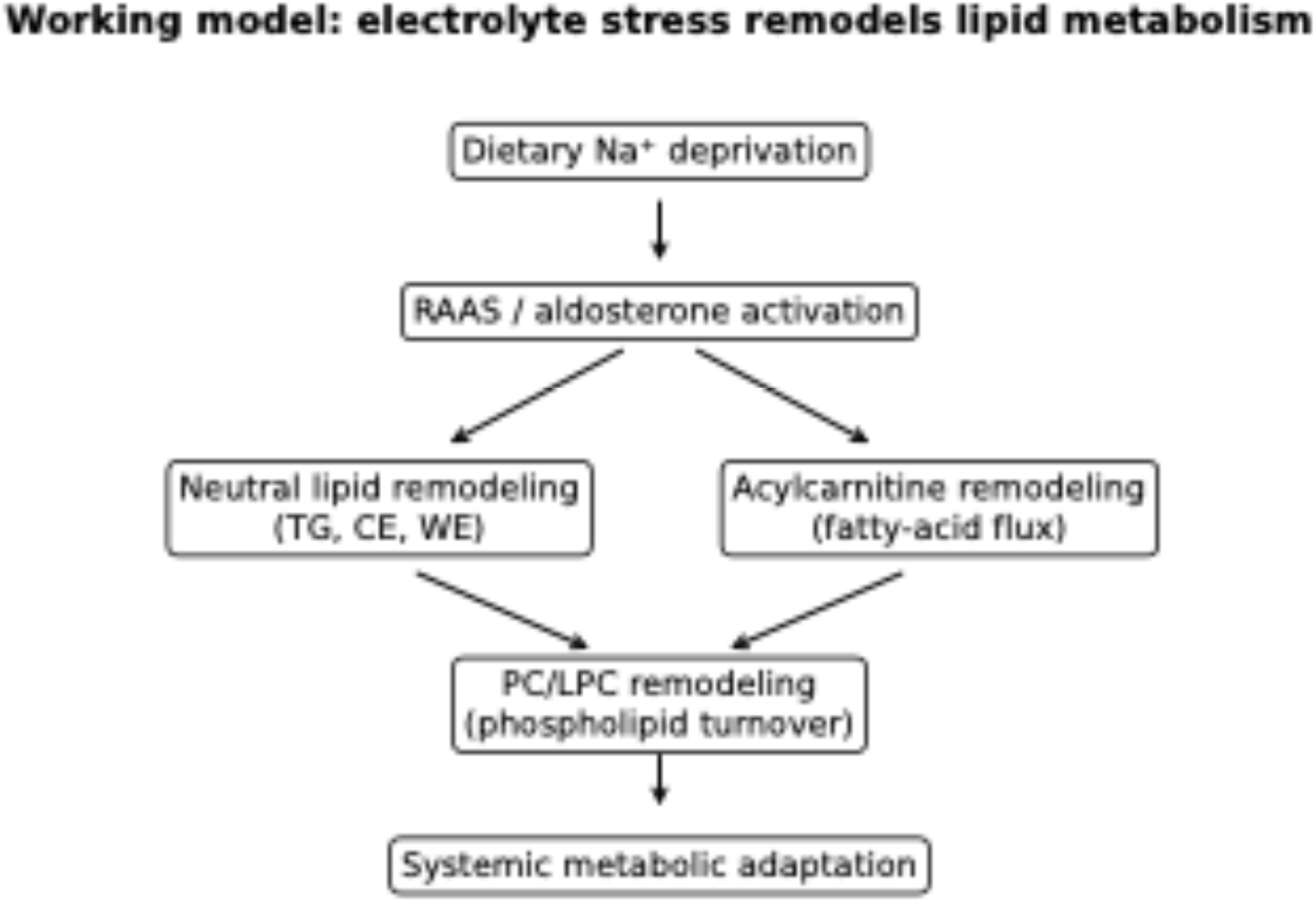
Working model: dietary sodium deprivation remodels circulating lipid metabolism. Schematic summarizing an integrative model derived from the serum lipidomics data. Dietary sodium deprivation is proposed to activate endocrine electrolyte-stress pathways (e.g., RAAS/aldosterone), leading to coordinated remodeling of neutral lipid storage pools (TG, CE, WE) and increased acylcarnitine-associated fatty-acid flux. These changes converge on enhanced membrane phospholipid turnover (PC/LPC remodeling) and collectively support a systemic metabolic adaptation to sodium restriction. This model is intended as a conceptual framework integrating the major lipid class signatures observed in the study.

This study has several limitations. First, the lipidomics analyses were performed in a small exploratory cohort (n=3 per group) after a short-term 7-day dietary intervention, which limits statistical power and generalizability. Second, the untargeted measurements are based on relative peak intensities and do not provide absolute lipid concentrations or direct metabolic flux information. Third, the analyses were restricted to serum and therefore do not identify the tissue sources driving the observed lipid remodeling. Finally, although the lipidomic signatures are consistent with altered lipid mobilization, phospholipid turnover, and mitochondrial fatty-acid handling, this study did not include direct mechanistic measurements such as tracer-based flux assays, tissue respiration assays, or endocrine profiling. These mechanistic interpretations should therefore be viewed as hypothesis-generating and warrant targeted validation in future studies of diet-electrolyte-metabolism interactions.

## CONCLUSIONS

Dietary sodium deprivation drives coordinated serum lipidome remodeling that reflects systemic metabolic adaptation at the level of neutral lipid mobilization, membrane organization, and mitochondrial fatty-acid handling. These findings support dietary sodium availability as a nutritional variable capable of shaping lipid metabolic phenotypes.

## ABBREVIATIONS

Na⁺: sodium
NaS: sodium-sufficient
NaD: sodium-deprived
TG: triglycerides
CE: cholesteryl esters
WE: wax esters
PC: phosphatidylcholine
LPC: lysophosphatidylcholine
AcCa: acylcarnitines
LC–MS/MS: liquid chromatography–tandem mass spectrometry
PCA: principal component analysis
FDR: false discovery rate
RAAS: renin–angiotensin–aldosterone system
SEM: standard error of the mean
VIP: variable importance in projection

## DECLARATIONS

## Authors’ contributions

J.C.H. conceived the study, contributed to data analysis and interpretation, and critically reviewed and edited the manuscript. S.K. performed experiments and contributed to data analysis. J.V. interpreted data and edited the manuscript. J.T.K. interpreted data and provided critical input. G.I.S. provided critical intellectual input, helped interpret the results, and edited the manuscript. V.M.R. conceived the study, performed experiments, supervised the project, and critically reviewed and edited the manuscript with input from all authors. All authors reviewed and approved the final version of the manuscript.

## Availability of data and materials

The processed lipidomics feature tables are provided as Supplementary Tables ST1 and ST2. Additional datasets generated and/or analyzed during the current study are available from the corresponding author upon reasonable request.

## Ethics approval and consent to participate

All animal experimental procedures used in this study were approved by the West Virginia University Institutional Animal Care and Use Committee prior to the start of the project. Consent to participate is not applicable.

## Consent for publication

Not applicable.

## Acknowledgements

Not applicable.

---

**Supplementary Figure S1.**
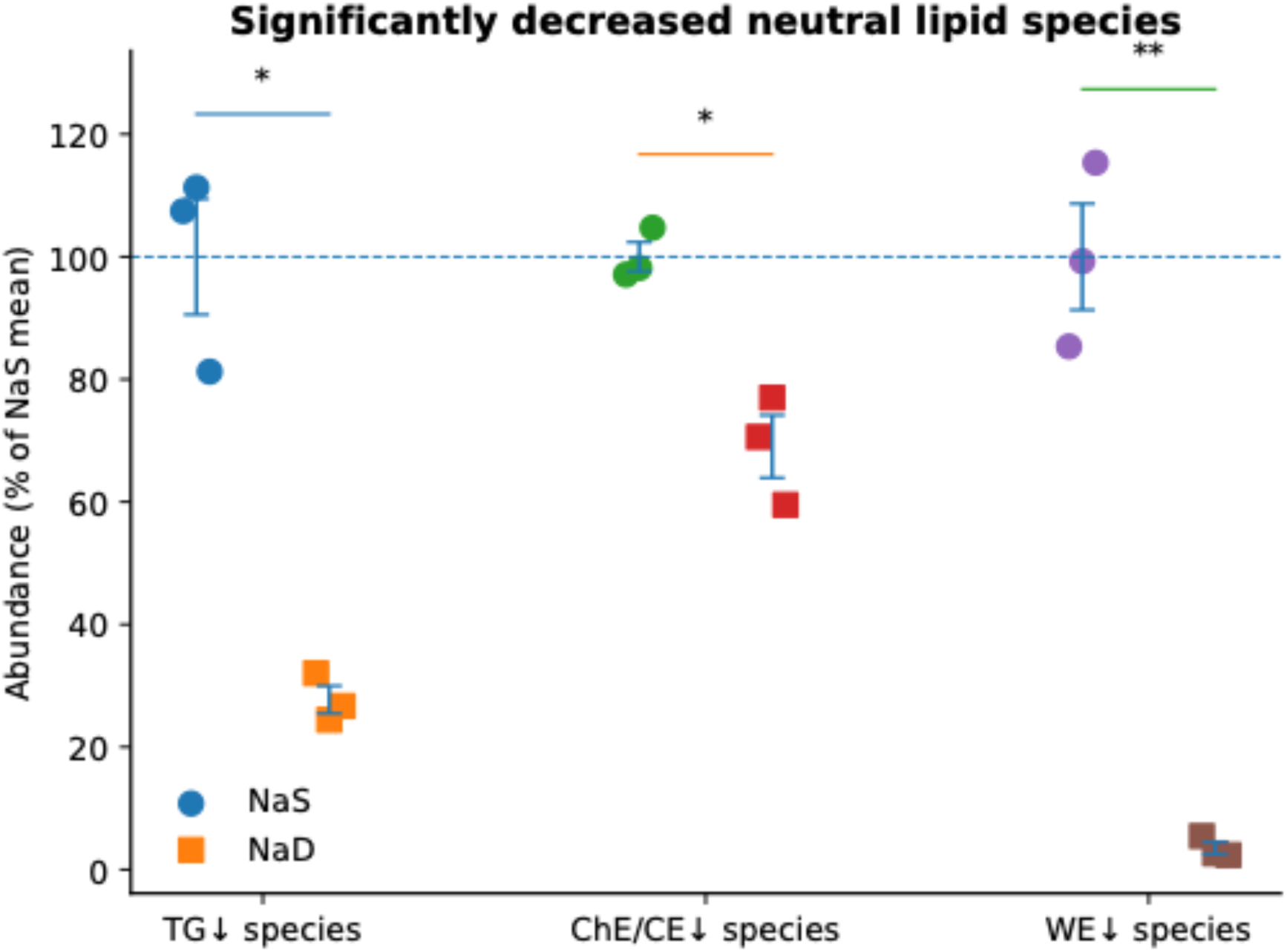
Dietary sodium deprivation markedly reduces selected neutral lipid storage species in serum. Scatter plot showing abundances of representative triglyceride (TG), cholesteryl ester (CE), and wax ester (WE) species in serum from sodium-sufficient (NaS) and sodium-deprived (NaD) rats. Values are expressed as percent of the NaS group mean (NaS=100%, dashed line) and plotted as individual biological replicates (n=3 per group) with mean ± SEM. Species displayed represent the subset of neutral lipid features identified as significantly decreased under sodium deprivation. Statistical significance was assessed using a two-tailed unpaired t-test for each species; P<0.05 (*), P<0.01 (**).

## REFERENCES

1. Noda M, Matsuda T (2022) Central regulation of body fluid homeostasis. Proc Jpn Acad Ser B Phys Biol Sci 98:283–324. 10.2183/pjab.98.016

2. Bernal A, Zafra MA, Simon MJ, Mahia J (2023) Sodium homeostasis, a balance necessary for life. Nutrients 15:395. 10.3390/nu15020395

3. Kaur J, Rout P (2026) Physiology, renin angiotensin system. In: StatPearls. StatPearls Publishing, Treasure Island (FL)

4. de Borst MH, Navis G (2016) Sodium intake, RAAS-blockade and progressive renal disease. Pharmacol Res 107:344–351. 10.1016/j.phrs.2016.03.037

5. Yoon H, Shaw JL, Haigis MC, Greka A (2021) Lipid metabolism in sickness and in health: emerging regulators of lipotoxicity. Mol Cell 81:3708–3730. 10.1016/j.molcel.2021.08.027

6. Jarc E, Petan T (2019) Lipid droplets and the management of cellular stress. Yale J Biol Med 92:435–452

7. Mahmoud AM, Mirza I, Metwally E et al (2025) Lipidomic profiling of human adiposomes identifies specific lipid shifts linked to obesity and cardiometabolic risk. JCI Insight 10. 10.1172/jci.insight.191872

8. Anwar MY, Highland HM, Palmer AB et al (2025) The circulating lipidome in severe obesity. medRxiv. 10.1101/2025.06.11.25329456

9. Surendran A, Zhang H, Stamenkovic A, Ravandi A (2025) Lipidomics and cardiovascular disease. Biochim Biophys Acta Mol Basis Dis 1871:167806. 10.1016/j.bbadis.2025.167806

10. Xu J, Lu D, Huang X (2026) Lipid metabolism and metabolites: emerging roles in systemic physiology and metabolic diseases. J Genet Genomics. 10.1016/j.jgg.2026.01.006

11. Markgraf DF, Al-Hasani H, Lehr S (2016) Lipidomics—reshaping the analysis and perception of type 2 diabetes. Int J Mol Sci 17:1841. 10.3390/ijms17111841

12. Callaghan NI, Durland LJ, Ireland RG et al (2022) Harnessing conserved signaling and metabolic pathways to enhance the maturation of functional engineered tissues. NPJ Regen Med 7:44. 10.1038/s41536-022-00246-3

13. Tao Z, Cheng Z (2023) Hormonal regulation of metabolism—recent lessons learned from insulin and estrogen. Clin Sci (Lond) 137:415–434. 10.1042/CS20210519

14. Briet M, Schiffrin EL (2011) The role of aldosterone in the metabolic syndrome. Curr Hypertens Rep 13:163–172. 10.1007/s11906-011-0182-2

15. Geisberger S, Bartolomaeus H, Neubert P et al (2021) Salt transiently inhibits mitochondrial energetics in mononuclear phagocytes. Circulation 144:144–158. 10.1161/CIRCULATIONAHA.120.052788

16. Tsai CH, Chen ZW, Lee BC, et al (2024) Aldosterone, mitochondria and regulation of cardiovascular metabolic disease. J Endocrinol 263. 10.1530/JOE-23-0350

17. Santos AL, Preta G (2018) Lipids in the cell: organisation regulates function. Cell Mol Life Sci 75:1909–1927. 10.1007/s00018-018-2765-4

18. Horn A, Jaiswal JK (2019) Structural and signaling role of lipids in plasma membrane repair. Curr Top Membr 84:67–98. 10.1016/bs.ctm.2019.07.001

19. Longo N, Frigeni M, Pasquali M (2016) Carnitine transport and fatty acid oxidation. Biochim Biophys Acta 1863:2422–2435. 10.1016/j.bbamcr.2016.01.023

20. Makrecka-Kuka M, Sevostjanovs E, Vilks K et al (2017) Plasma acylcarnitine concentrations reflect the acylcarnitine profile in cardiac tissues. Sci Rep 7:17528. 10.1038/s41598-017-17797-x

21. Wang B, Tontonoz P (2019) Phospholipid remodeling in physiology and disease. Annu Rev Physiol 81:165–188. 10.1146/annurev-physiol-020518-114444

22. Zhang Q, Cheng L, Li H et al (2021) The structural basis for the phospholipid remodeling by lysophosphatidylcholine acyltransferase 3. Nat Commun 12:6869. 10.1038/s41467-021-27244-1

23. Nicolson GL, Ferreira de Mattos G, Ash M, Settineri R, Escriba PV (2021) Fundamentals of membrane lipid replacement. Membranes (Basel) 11:944. 10.3390/membranes11120944

24. Cesari AB, Paulucci NS, Dardanelli MS (2026) Adaptive physiological and membrane lipid responses of Bradyrhizobium sp. SEMIA 6144 to osmotic stress. J Appl Microbiol 137. 10.1093/jambio/lxag088

25. Du M, Chen J, Wang C, et al (2025) Membrane lipid remodeling strategies regulate fluidity for acute temperature adaptation in oysters. Evol Appl 18:e70156. 10.1111/eva.70156

26. Barigou M, Ramzan I, Chartoumpekis DV (2025) The role of aldosterone and the mineralocorticoid receptor in metabolic dysfunction-associated steatotic liver disease. Biomedicines 13:1792. 10.3390/biomedicines13081792

